# Dynamical model of the CLC-2 ion channel reveals conformational changes associated with selectivity-filter gating

**DOI:** 10.1101/228163

**Authors:** Keri A. McKiernan, Anna K. Koster, Merritt Maduke, Vijay S. Pande

## Abstract

This work reports a dynamical Markov state model of CLC-2 “fast” (pore) gating, based on 600 microseconds of molecular dynamics (MD) simulation. In the starting conformation of our CLC-2 model, both outer and inner channel gates are closed. The first conformational change in our dataset involves rotation of the inner-gate backbone along residues S168-G169-I170. This change is strikingly similar to that observed in the cryo-EM structure of the bovine CLC-K channel, though the volume of the intracellular (inner) region of the ion conduction pathway is further expanded in our model. From this state (inner gate open and outer gate closed), two additional states are observed, each involving a unique rotameric flip of the outer-gate residue GLU_*ex*_. Both additional states involve conformational changes that orient GLU_*ex*_ away from the extracellular (outer) region of the ion conduction pathway. In the first additional state, the rotameric flip of GLU_*ex*_ results in an open, or near-open, channel pore. The equilibrium population of this state is low (∼1%), consistent with the low open probability of CLC-2 observed experimentally in the absence of a membrane potential stimulus (0 mV). In the second additional state, GLU_*ex*_ rotates to occlude the channel pore. This state, which has a low equilibrium population (∼1%), is only accessible when GLU_*ex*_ is protonated. Together, these pathways model the opening of both an inner and outer gate within the CLC-2 selectivity filter, as a function of GLU_*ex*_ protonation. Collectively, our findings are consistent with published experimental analyses of CLC-2 gating and provide a high-resolution structural model to guide future investigations.

**Author summary:** In contrast to sodium-, potassium-, and calcium-selective ion channels, the roles of chloride-selective ion channels in mammalian physiology have been studied much less and are not sufficiently understood, despite known associations of chloride-channel defects with a variety of pathological conditions. CLC-2 is a voltage-activated chloride channel (one of 9 human CLC homologs) with broad tissue and organ distribution. In this work, we use simulations to model the conformational dynamics of the CLC-2 chloride ion channel selectivity filter (SF), which is the part of the protein that controls whether the channel is in an ion-conducting or non-conducting state. Our analysis identifies four primary conformational states and a specific progression through these states. Our results are consistent with structural and functional data in the literature and provide a high-resolution model for guiding further studies of CLC-2. These results will inform our understanding of how CLC-2 regulates electrical activity and ion homeostasis in the many tissues where it is expressed.

## Introduction

The CLC family plays a wide variety of physiological functions in organisms ranging from bacteria to humans [1–7]. This family of membrane proteins is composed of both channels and H^+^/Cl^−^ exchange transporters that share a structurally unique homodimeric architecture [8, 9]. Each subunit within the homodimer is an independent functional unit [10–12] composed of 17 membrane-embedded alpha helices [13]. These helices coalesce to form a narrow, electropositive ion-conducting pore that is highly selective for Cl^−^ [13]. Such an architecture is unusual for ion channels and raises many questions about the mechanisms of ion channel opening and closing (gating).

High-resolution CLC structures provide invaluable starting points for understanding the structure-function relationships of CLC protein activation. The first CLC structure solved, a prokaryotic H^+^/Cl^−^ exchange transporter [13], proved relevant for guiding structure-function studies on a wide range of CLCs, including eukaryotic CLC channels [15, 16, 18, 19]. The mechanistic similarities between channels and transporters, which have been discussed and studied extensively, provide strong justification for using CLC transporter structures to understand CLC channel structure and function [8, 20]. The CLC transporter structures represent a closed conformational state in which bound chloride (Cl^−^) is occluded by proteinaceous “gates” that block exit to either the outer (extracellular) or inner (intracellular) side of the protein. The outer gate is formed in part by a conserved glutamate residue, GLU_*ex*_ [21–28] (Fig. 1). When mutated to GLN (to resemble the protonated GLU), this side chain rotates outwards and partially unblocks the Cl^−^ permeation pathway [21]. The inner gate is formed by conserved SER (SER_*cen*_) and TYR residues [13, 29–31] that physically obstruct the Cl^−^ translocation pathway from the intracellular side.

**Fig 1.**
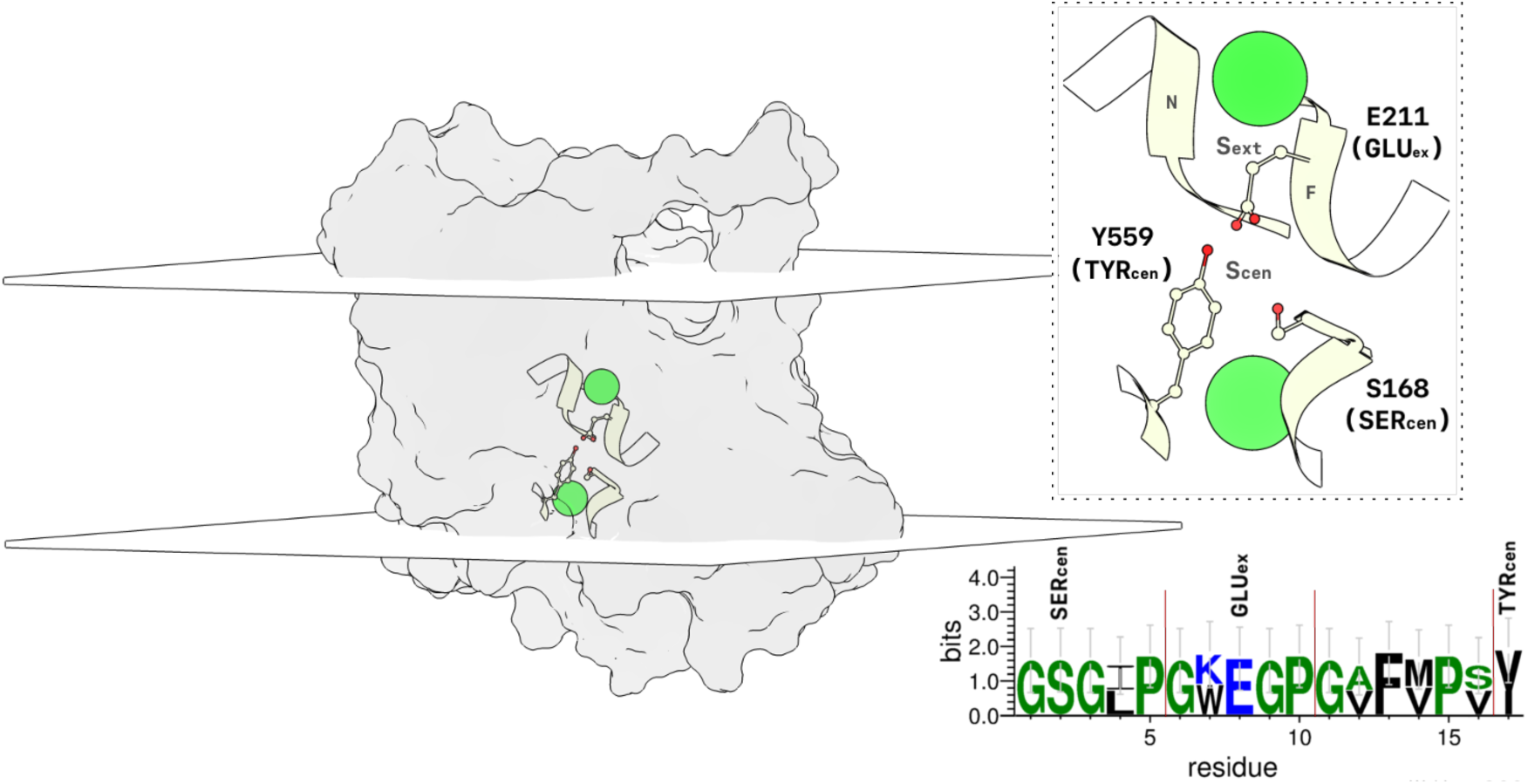
Illustration of the rCLC-2 monomer simulated in this study. The protein is shown as a transparent surface. The planar surfaces represent the system membrane bilayer (planes intersect the average z-coordinate of the lipid phosphorus atoms). The SF is highlighted in a ribbon representation. A larger view of the SF is given by the inset with a dashed border. Mechanistically important Cl^−^ binding sites, S_*cen*_ and S_*ext*_, are labeled, as are nearby residues GLU_*ex*_, SER_*cen*_, and TYR_*cen*_. Helices F and N help form the S_*ext*_ binding site. GLU_*ex*_, on Helix F, is positioned at the S_*cen*_ ion binding site in this conformation. Chloride ions are shown in green. Sequence conservation between the CLC-2 and cmCLC (3ORG) SF residues is displayed on the bottom right. Hydrophilic residues are highlighted in blue, neutral residues in green, and hydrophobic residues in black. Residue height denotes the relative frequency of the amino acid at a given position. When there is a residue discrepancy, the CLC-2 residue is given on top. Each region of the SF has been segmented by the red lines.

Following the completion of the simulation studies described herein, the structures of two mammalian CLC channels were published, bovine CLC-K (bCLC-K) and human CLC-1 (hCLC-1) [32, 33]. These structures are highly similar to the transporter structures, but with subtle structural differences that result in wider anion permeation pathways at both the extracellular and intracellular gates. In the bCLC-K structure, widening at the extracellular gate is due to the replacement of GLU_*ex*_ with a neutral VAL residue. In CLC-1, widening at the extracellular pore entrance instead results from outward rotation of GLU_*ex*_. At the intracellular side, widening of the Cl^−^ permeation pathway is accomplished in one of two ways. In the bCLC-K structure, the loop containing SER_*cen*_ undergoes a conformational change such that the SER_*cen*_ side chain rotates to unblock the pore. In the hCLC-1 structure, no rotation of SER_*cen*_ occurs. Rather, expansion of the intracellular pore entrance results from the smaller volume of residues in this region. While these new structures provide invaluable molecular detail for understanding the gating mechanisms of CLC channel homologs compared to their transporter counterparts, many questions remain. What, for example, is the conformation of the CLC-K closed state, given that it does not have a GLU_*ex*_ gate? Does the intracellular conformational change seen in CLC-K (the rotation of the conserved SER-containing loop) occur also in CLC-1 (and CLC-2) to further widen the pore? How does the hyperpolarization-activated open state of the CLC-2 channel compare to the open states of CLC-K (non-voltage dependent) and CLC-1 (depolarization-activated)? Answering these questions will require turning these static conformational snapshots into a complete dynamic picture of CLC gating. Experimental biophysicists have shown that these mechanisms are complex, involving interdependent influences of transmembrane voltage, [Cl^−^], and [H^+^] [8, 34–38]. To begin to resolve these complexities and reveal the dynamic, atomistic details of the CLC gating machine, it will be critical to pair structural studies with high-resolution conformational modeling [39–48].

In this study, we investigate the molecular dynamics (MD) of CLC-2, a mammalian chloride ion channel with broad tissue and organ distribution [49, 108]. The widespread physiological importance of CLC-2 is reflected by the severe phenotype of knockout animals, which exhibit male infertility, blindness, leukodystrophy, and decreased intestinal Cl^−^ absorption [113–115]. In humans, CLC-2 defects are linked to leukodystrophies [108], and gain-of-function mutations result in primary aldosteronism [116]. Despite the links of CLC-2 malfunction to human disease, many aspects of CLC-2 physiology remain uncertain. For instance, while CLC-2 is the most broadly expressed voltage-gated chloride channel in the brain, the details of its role in neuronal excitability and/or chloride homeostasis [55–63] are controversial [3, 62, 64, 65]. A precise understanding of the atomic interactions and rearrangements involved in channel activation will facilitate understanding of CLC-2 physiological behavior and its role in human disease.

The spatial and temporal resolution afforded by computer simulations allows for the atom-scale examination of protein structure and function. Here we apply MD simulations and Markov State Modeling to study the conformational dynamics of CLC-2 gating. Markov State Models (MSMs) allow for the rigorous statistical characterization of biophysical simulations [66–69]. Prior to constructing the MSM, the simulation data is first transformed into its kinetically slowest representation via the method of time-structure based independent component analysis (tICA) [70–72]. Such a transformation allows identification of the slowest conformational changes in an MD dataset - those most likely to correspond to channel gating. The MSM is then built over these conformational changes, and queried to predict the thermodynamics and kinetics of CLC-2 channel activation. Our MSM indicates that the top activation pathway for our CLC-2 model begins in the closed state, near the template structure on which we based our model. The slowest timescale conformational change involves the rotation of the backbone along the S168-G169-I170 residues at the intracellular entrance to the ion conduction pathway. The second major motion describes the flip of the GLU_*ex*_ side chain on the extracellular vestibule. This flip of the GLU_*ex*_ side chain displays a sensitivity to residue protonation that correlates well with experimental evidence that this conserved glutamate residue regulates the CLC “fast” gate in CLC-2 [73, 74]. These observed gating motions are consistent with data in the literature. This high-resolution structural model of CLC-2 gating will facilitate experimental design and analysis of future mechanistic studies involving CLC channels.

## Results and discussion

Ion channel “gating” involves the conformational rearrangement of the protein, such that transmembrane ion flow through the channel pore is either increased or decreased. These gates are highly regulated so as to avoid dissipation of cellular ion gradients. CLC channels have two types of gating mechanisms, known as slow and fast gating. Slow gating (also known as “common gating”) involves activation of both subunits of the homodimer in unison, requiring global conformational changes and communication between subunits at the dimer interface [12, 75–77]. Fast gating (also known as “pore gating”) acts on each subunit independently and involves much smaller conformational changes, perhaps limited to motions of side chains within the selectivity filter (SF) [8, 9, 78]. Studies based on CLC transporter structures demonstrate similarities between CLC channel fast gating and the “gating” involved in CLC transporter activity. Specifically, high-resolution structures of CLC transporters reveal that a conserved glutamate residue (GLU_*ex*_) physically occludes the permeation pathway, blocking Cl^−^ access from the extracellular side. In transporters, mutation of this residue to glutamine causes the side chain to swing upward, permitting an additional Cl^−^ ion to occupy the pore [21, 24–26]. In channels, mutation of GLU_*ex*_ to a neutral equivalent similarly unblocks the permeation pathway, as revealed by the fast-gate open phenotype that is observed by electrophysiological recordings [21, 22, 28]. Thus, CLC channel fast gating is highly analogous to CLC transporter gating and involves movement of GLU_*ex*_ away from the extracellular Cl^−^ binding site (S_*ext*_).

Despite the understanding gained from these combined structural and electrophysiological analyses, questions remain concerning CLC fast gating. For example, are additional conformational changes at the extracellular side involved in channel opening, as has been suggested to occur for the transporters [41]? Further, is there an intracellular gate involved in fast gating? While the structures of CLC-K and CLC-1 suggest that the inner gates may be wide enough to support Cl^−^ permeation, electrophysiological studies have suggested that CLC pore gating involves movement of inner-gate residues. This idea is based on studies examining altered state-dependent binding of inhibitors at the intracellular side of a CLC channel. These studies concluded that such behavior could only be explained by conformational changes occurring at the inner gate, in conjunction with the side chain rotation of GLU_*ex*_ at the outer gate [79]. Using the CLC transporter structures as a guide, these inner-gate residues correspond to conserved SER_*cen*_ and TYR_*cen*_ residues that physically prevent Cl^−^ from entering the pore [13, 27, 29–31]. Consistent with the conclusion that these residues must move in order to open the channel, the recent CLC-K channel structures show that these residues adopt a more open configuration that results from rotation of a loop containing the intracellular SER_*cen*_ gate. Intriguingly, this rotation is not observed in the CLC-1 structure, suggesting that such a change may not be needed to open that CLC channel homolog. Additional work is necessary to determine the dynamic details of how conformational changes at the inner and outer gates are linked to opening CLC channel pores. Because CLC channel homologs differ in the degree to which the pore is gated by Cl^−^, H^+^, transmembrane voltage, and other physiological variables [73, 74, 80–86], the details of pore gating are likely to vary subtly between the homologs, despite the overall similarity in their structures. In this work, we sought specifically to gain insight into pore behavior in the CLC-2 homolog.

We applied dynamical modeling in order to identify and piece together the major conformational changes characterizing inner- and outer-gate opening in CLC-2. First, a homology model was built for the rat CLC-2 (rCLC-2) sequence, using the 3ORG structure of the cmCLC transporter [87]. This template was chosen because, at the time of construction, it was the only available eukaryotic CLC structure. This template is 75% identical to CLC-2 within the pore-forming (SF) regions of the protein. The sequence conservation between the target and template [88, 89] SF is depicted in Fig. 1, bottom right, with conserved outer- and inner-gate residues (E211, Y559, and S168) depicted above. The template and target sequences have a global similarity of 34.4%. Notably, transmembrane regions with sequence similarity in this range have been shown to display high structural similarity [90]. Stability of the homology model was evaluated via Ramachandran plot statistics [112]. Only 0.2% of residues in the model were found to exist in disallowed regions, this is compared to the template crystal structure which was found to have 0.3% of residues in disallowed regions. Additional statistics are given in **??**. From this model, two MD datasets were generated, identical with the exception of GLU_*ex*_ protonation. These datasets were analyzed with respect to the SF conformation (Fig. 1 inset). To improve computational sampling, two simplifications were made in our structural model. First, because each subunit contains an independent and identical pore [10–12], only one subunit of a homodimer was simulated. This simplification is justified based on the fact that fast gating occurs independently within each subunit pore [10–12, 78]. Additional evidence for the existence of independently stable and functional CLC subunits comes from studies of purified, reconstituted CLC-ec1 protein. The free energy of WT CLC-ec1 dimerization in bilayers is measurable [109], and functional CLC-ec1 monomers can be generated by a simple double mutation that disrupts interactions at the dimer interface [111]. In accordance with these experimental observations, the CLC-2 monomer was found to be stable in our MD simulation. The second simplification in our model is that the C-terminal cytoplasmic domain was removed from the model. Removal of this domain has minimal effects on CLC-2 channel properties, resulting in nearly identical single-channel current and maximal open probability, relative to WT, with a slightly shifted voltage dependence of activation (by 10 mV). The open probability at 0 mV, where our studies are conducted, are indistinguishable between WT and C-terminal-deleted CLC-2. The biggest effect of C-terminal deletion is on channel activation and deactivation rates, with the deletion mutant showing rates about five-fold faster than WT CLC-2 [107]. Our model also lacks the N-terminal residues 1-83, which are not present in the original cmCLC template. Similar to removal of the C-terminal domain, partial deletion of the N-terminus (Δ16-61) exhibits a voltage-dependent open probability identical to WT CLC-2, though with faster activation and deactivation rates [110]. These simplifications are thus experimentally justified and are also advantageous from a computational point of view, reducing the number of atoms in the simulation and the amount of sampling required to observe interesting conformational changes. While the kinetics of transitions between the states in our truncated computational model will likely differ from WT, experimental evidence is strong that the overall CLC-2 fast-gating machinery remains intact for our simplified CLC-2 construct. Although we did not explicitly account for specific Cl^−^/protein interactions in our model, the effects of Cl^−^ on channel gating are implicitly considered by its presence in our simulation, providing a kinetic framework for future investigations into the specific mechanisms of ion permeation. The CLC-2 construct simulated in this study is depicted in Fig. 1.

### Selectivity filter conformational dynamics

Markov State Models have emerged as a powerful tool for robustly estimating the statistics of biophysical systems from MD datasets [66–69]. This technique first partitions a dataset into a set of discrete conformational states, and then parameterizes a transition matrix describing the probability of transition for each state pair. Prior to constructing the MSM, it is advantageous to featurize the data according to a structural metric, and then transform the featurized dataset using time-structure independent component analysis (tICA) [70]. tICA is very good at describing both local motions and global motions, but not at the same time given the very different timescales of these processes. We chose to focus on local motions because published data all strongly point to such motions as playing a key role in CLC channel fast gating [8, 9, 78]. Although this approach will necessarily miss larger-scale conformational change that may be involved in fast gating [41], it will discern fine details of local conformational intermediates, which are critical to understanding CLC-2 gating. An additional advantage of this approach is that we have very high confidence in our model in this region of the protein, given the very high sequence conservation (71% similar, 88% identical between rCLC-2 and the cmCLC template – **??**). We did attempt to construct the model from the perspective of macroscopic whole-protein conformation change, but as expected, it was not as effective at identifying all selectivity filter states. (The low-probability *O* and *U* states described below cannot be detected with this approach.)

To focus on local conformational change, we used the backbone and side chain dihedral angles of 17 SF residues as the structural metric for featurization (see the Markov State Modeling subsection of the Methods section for more detail). tICA determines the slowest-decorrelating dynamical degrees of freedom [91]. This improves the MSM statistics, and additionally provides a set of unbiased reaction coordinates that are useful for separating and visualizing important conformational states. These reaction coordinates can be thought of as dynamical modes, and are referred to as “tICs”. A tIC is given by a linear combination of input features, and can be interpreted by identifying the features with the highest linear coefficients (referred to as loadings). Greater detail describing the tIC loadings is given in **??**. After the MSM is constructed, the state representation can be simplified via macrostating. This simplification allows for the identification of the most thermodynamically stable conformational states. Here, we employ these techniques to characterize the conformational dynamics sampled in our datasets.

Our analysis identified four major conformational states: *C*_*oi*_, *C*_*o*_, *O*, and *U*. These states were named to reflect whether the channel appears closed (C) or open (O) at the inner (i) or outer (o) gate, with the exception of *U*, a postulated non-conducting state. A representative conformation for each thermodynamic macrostate is illustrated in Fig. 2. The nomenclature of these states represents the x, y, z-positions of the inner and outer gating residues identified in the MSM analysis, as well as our analysis of the pore radius for each macrostate (see Fig. 5). As such, the names of these four states do not necessarily represent the entire collection of open and closed states possible under physiological conditions, but rather whether the entrance and exit to the pore are physically blocked and the extent to which each conformation can accommodate the volume of a Cl^−^ ion. It is possible that other conformational states (such as a more fully open conducting state) were not sampled in our 600 microsecond experimental timescale due to the extremely long (seconds) timescale required for full CLC-2 channel activation, as well as the limitations imposed by modeling the channel at 0 mV instead of hyperpolarizing potentials (at which the channel has the highest open probability). Despite these limitations, the conformational states that we observe involve motions of SF residues that unblock the pore and widen the anion permeation pathway, suggesting a physiologically relevant mechanism of channel activation. These states are also highly consistent with what is experimentally known about the fast-gate mechanism of CLC channel opening (subsequently discussed in detail). For a detailed labeling of each SF component, see Fig. 1.

**Fig 2.**
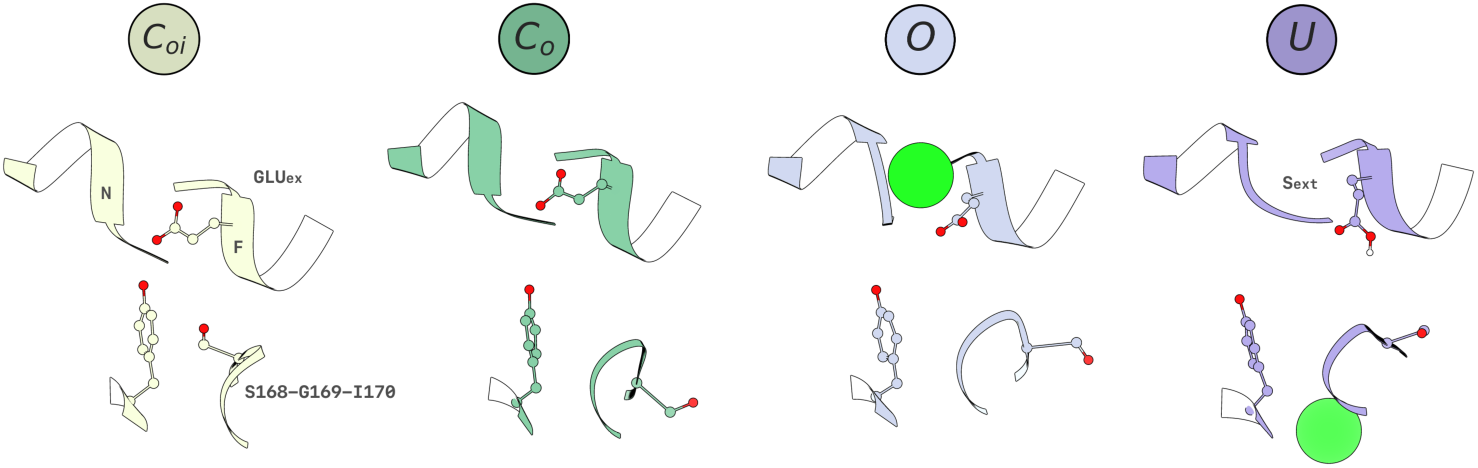
Structural visualization of each CLC-2 SF macrostate conformation. The S168-G169-I170 residues, located at the inner region of the SF, exist in two orientations, either pointing toward the ion conduction pathway (*C*_*oi*_), or away (*C*_*o*_, *O, U*). The GLU_*ex*_ residue side chain, located at the outer region of the SF, exists in three orientations, either occluding the S_*ext*_ Cl^−^ binding site, located in the gap between helices F and N (*C*_*oi*_, *C*_*o*_), or in one of the two distinct outward rotamers (*O* and *U*). Note that in the *U* state, the geometry of the S_*ext*_ binding site is distorted by the conformation of helix N.

Each macrostate can be kinetically distinguished from the others by tIC 0, tIC 1, and tIC 3 (tIC 2 describes the same conformational change as tIC 1 and hence will not be discussed; such similarity can result from symmetries of the featurized data). tICs 1-3 were interpreted by analyzing the input features with the highest loadings. In Table 1 we have detailed how each tIC (dynamical mode) separates a particular two-state kinetic process, and which features distinguish each tIC.

**Table 1.** Dynamical modes of the MD data, as determined by tICA.

| dynamical mode | conformational change | top residue features | kinetic process |
| --- | --- | --- | --- |
| tIC 0 | SER<sub>cen</sub> backbone rotation | $\psi(\text{I170,G169})$, $\psi(\text{G169,S168})$ | $C_{oi} \leftrightarrow C_o$ |
| tIC 1 | GLU<sub>ex</sub><sup>p</sup> side chain flip | $\chi_1(\text{E211})$, $\phi(\text{S168,G167})$ | $C_o \leftrightarrow U$ |
| tIC 3 | GLU<sub>ex</sub> side chain flip | $\chi_1(\text{E211})$, $\phi(\text{G169,S168})$ | $C_o \leftrightarrow O$ |

Along each major dynamical mode, we observe two-state behavior. tIC 0 involves the rotation of the backbone dihedrals of the S168, G169, and I170 residues. This implies that the bottom strand of the SF can exist in two unique conformations. tICs 1 and 3 both involve flipping the GLU_*ex*_ *χ*_1_ side chain dihedral, a motion that is coupled to the residues involved in the tIC 0 conformational change. Although tICs 1 and 3 begin in the same initial state (*C*_*o*_) and describe the rotameric flipping of the same residue, the end states (*O* and *U*) differ in conformation. This suggests that the GLU_*ex*_ rotamer can exist in three possible states. ?

The thermodynamics of each conformational state was further analyzed using the MSM-derived free energies. This analysis is illustrated by Fig. 3, where the data have been separated by the GLU_*ex*_ protonation state, and projected onto each major dynamical mode. We observe that in all cases, *C*_*oi*_ describes the conformational free energy minimum. This state, so named because the channel pore appears closed to both the outer (o) and inner (i) sides of the membrane, is similar to the template cmCLC structure (3ORG) with the exception that GLU_*ex*_ has moved upwards out of the S_*cen*_ site, towards the S_*ext*_ site, in a position similar to that observed in most other CLC structures found in the PDB. The *C*_*o*_ state is the next most stable state, with a free energy less than 1 kCal/mol greater than the *C*_*oi*_ state. This state was labeled *C*_*o*_ because the channel pore is closed to the outer side but open towards the inner side. In this state, the S168, G169, and I170 backbone strand has changed conformation, but the GLU_*ex*_ residue remains in the same geometry. As can be seen by the landscapes in Fig. 3, GLU_*ex*_ protonation does not affect the free energies of the *C*_*oi*_ and *C*_*o*_ states. The transition barrier height for the *C*_*oi*_ ↔ *C*_*o*_ conformational change is approximately 2 kCal/mol.

**Fig 3.**
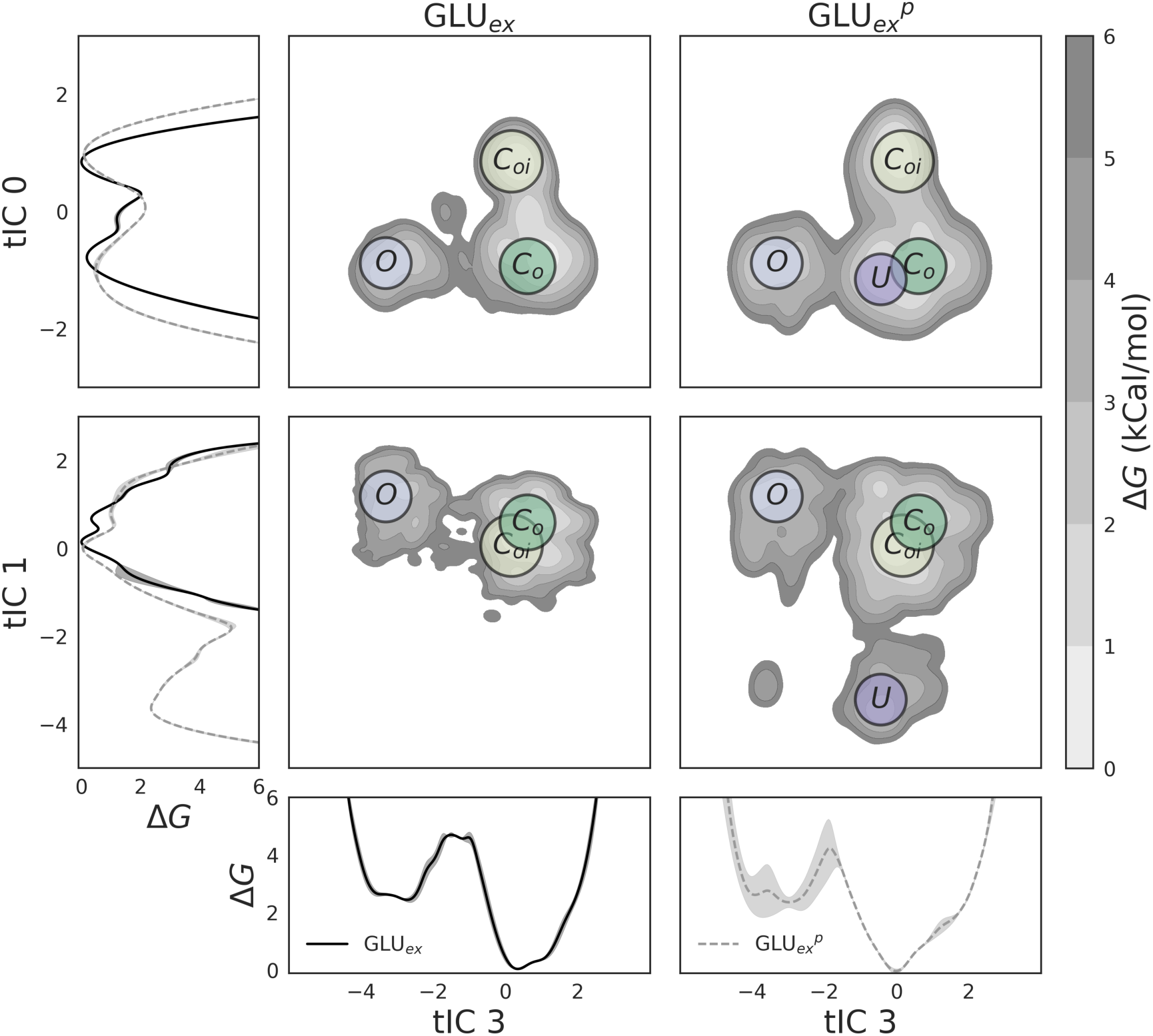
Free energy of major CLC-2 conformational states, projected onto all major dynamical modes (tICs), as a function of GLU_*ex*_ protonation. These landscapes are shown in both 1D and 2D, for both the deprotonated dataset (GLU_*ex*_, black solid lines) and the protonated dataset (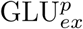, gray dashed lines). tIC 0 denotes the conformation of the S168-G169-I170 backbone, while tIC 1 and tIC 3 denote the conformation of the GLU_*ex*_ side chain. In all cases, *C*_*oi*_ was found to be the global free energy minimum, and to not vary greatly with respect to GLU_*ex*_ protonation. Also insensitive to protonation are the basin free energies of *C*_*o*_ and *O*. The largest difference caused by GLU_*ex*_ protonation is the existence of *U*, which occurs only for 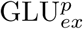 (as isolated by tIC 1), and the structure of the kinetic barriers.

tICs 1 and 3 identify two higher energy conformational states, *O* and *U*, each connected to state *C*_*o*_. As discussed previously, these states each represent rotamers of the GLU_*ex*_ residue. In the *O* (“open”) state, GLU_*ex*_ appears to move out of the way of the Cl^−^ permeation pathway, whereas in the *U* state, GLU_*ex*_ collapses into the pore, which remains occluded (as discussed below). Both states *O* and *U* are between 2 and 3 kCal/mol greater in energy than states *C*_*o*_ and *C*_*oi*_. The *O* state shows a mild sensitivity to protonation, while the *U* state exists only when GLU_*ex*_ is protonated. This result can be seen by the absence of a *U* basin in the GLU_*ex*_ 2D free energy plots of Fig. 3. The transition barrier height of *C*_*o*_ → *O*, along tIC 3, is approximately 4-5 kCal/mol for both protonation states, though there is considerable uncertainty for 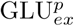 (shown by the error bar along tIC 3). The transition barrier height of *C*_*o*_ → *U*, along tIC 1, for GLU_*ex*_^*p*^ is approximately 5 kCal/mol.

Interestingly, TYR_*cen*_ (Y559) (Fig. 1) was not identified by any of the dynamical modes in our analysis. In prokaryotic CLC transporter structures and in our eukaryotic template structure (cmCLC), this residue, along with SER_*cen*_ (S168), forms a chloride-binding site in the pore. As such, mutation of these residues would be predicted to alter single-channel conductance by changing the binding energy of chloride in the pore. While mutagenesis experiments on SER_*cen*_ support this prediction, those on TYR_*cen*_ do not [11, 19, 79, 92]. In agreement with these findings, our model predicts movement of the SER_*cen*_ backbone (S168-G169-I170) to be critical for initiation of chloride translocation, while TYR_*cen*_ does not play a major role.

### Kinetic modeling and comparison to experimental data

To examine kinetics in greater detail, continuous-time rate-matrix MSMs were computed for the macrostate model in order to estimate the relative rate of transition between each major conformational state (as the traditional MSM gives transition probabilities, rather than transition rates). The resultant kinetic network graphs are given in Fig. 4. We observe that the *C*_*oi*_ state transitions only to the *C*_*o*_ state, while the *C*_*o*_ state may transition to any of the other states. The forward and backward rate of *C*_*oi*_ ↔ *C*_*o*_ are comparable. For both the *O* and *U* states, the reverse rates from these states to the *C*_*o*_ state are faster than the forward rates.

**Fig 4.**
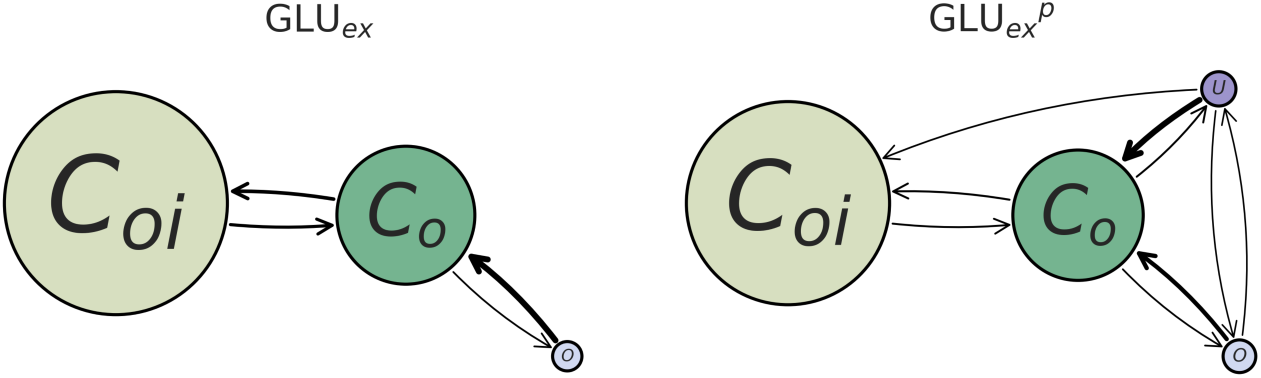
Kinetic rates describing the transition between major CLC-2 conformations, as a function of GLU_*ex*_ protonation. Note that the size of each node is proportional to its equilibrium population (larger denotes a lower free energy), and the weight of each arrow is proportional to the speed of the kinetic transition (heavier weight denotes a faster rate). Additionally, it is possible that there exist transitions occurring over timescales longer than what was sampled in our datasets (which likely underlies the lack of an observed transition from *C*_*oi*_ to *U*). As identified by the thermodynamic analysis, the *U* conformational state exists only for the 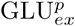 model. Both the GLU_*ex*_ and 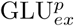 models show similar rates describing *C*_*oi*_ ↔ *C*_*o*_ and *C*_*o*_ ↔ *O*. However, in the case of 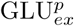, state *U* may transition forward to any other state.

The hypothesized non-conducting state is consistent with experiments demonstrating the state-dependent effect of pH on the CLC-2 ion channel. Under a constant hyperpolarizing stimulus of − 100 mV (to open the channel), application of pH 5.5 external buffer (to favor GLU_*ex*_ protonation) followed by a depolarizing pulse to close the channel (while GLU_*ex*_ remains protonated) renders the channel unable to re-open upon application of a second − 100 mV stimulus [93]. In contrast, the same experiment performed under basic conditions (pH 8.0, favoring GLU_*ex*_ deprotonation) reduces current overall but does not prevent the channel from re-opening, suggesting that GLU_*ex*_ protonation locks the channel in a unique non-conducting conformation that prevents immediate reactivation [93]. This observation is consistent with our model in which the non-conducting *U* conformational state of CLC-2 is only accessible when GLU_*ex*_ is protonated.

While the primary effect of GLU_*ex*_ protonation is the addition of the *U* state, kinetic analysis highlights two additional, mechanistically relevant, features of protonation. First, the connectivity of the modeled closed state, *C*_*oi*_, to the modeled open state, *O*, suggests that protonation of GLU_*ex*_ is not essential for channel conduction. While this result is in contrast to a computational study of the 3ORG (cmCLC) structure, which found that protonation of the GLU_*ex*_ side chain is required for ion transport to occur [42], it is in harmony with electrophysiological studies of CLC-2, which have demonstrated that protonation of GLU_*ex*_ does not affect channel opening [73]. The second additional highlight of our kinetic analysis is that the transition rate from *O* to *C*_*oi*_ is slowed when GLU_*ex*_ is protonated. While this result is uncertain due to the magnitude of the error bars of our calculations, it is worth highlighting given its consistency with experimental results, which demonstrate that the CLC-2 closing rate decreases with increasing extracellular [H^+^] [74].

### Structural analysis and comparison with experimental data

Above, we computed estimates for the thermodynamic and kinetic behavior of four major CLC-2 conformational states. Now, we delve deeper into the structural analysis of these states. To further investigate the geometric structure of each macrostate, the volume of the ion conduction pathway was analyzed. For each macrostate, several conformations were sampled, and the radius of the ion conduction pathway computed. These results are given with respect to distance from the GLU_*ex*_ residue (one path to the extracellular region, one path to the intracellular region), and are illustrated in Fig. 5. It was found that GLU_*ex*_ is the bottleneck of the conduction pathway for all macrostates. Of note, these bottlenecks are all significantly wider than the ∼ 0.2 Å bottleneck seen in the transporters [41], including the cmCLC transporter that was used as our template structure.

The three positions occupied by GLU_*ex*_ in CLC-2 are compared visually in Fig. 6A, right panel, which shows superpositions of representative conformations from macrostates *C*_*o*_, *O*, and *U* (*C*_*oi*_ and *C*_*o*_ have identical GLU_*ex*_ conformations). To compare these to experimental structures, we show relevant overlays in Fig. 6B. In the CLC transporter structures, the GLU_*ex*_ side chain has been observed to adopt three conformations, referred to as “down”, “middle”, and “up” [29]. The “down” conformation, with GLU_*ex*_ pointing into S_*cen*_, has been seen only in 3ORG (cmCLC), our template structure. In the CLC-2 model based on this structure, we found that GLU_*ex*_ immediately equilibrates away from this conformation to a rotamer where it occupies the S_*ext*_ binding site (the “middle” conformation), as in CLC-ec1. The GLU_*ex*_ “up” conformation, which has been observed when GLU_*ex*_ is mutated to GLN in CLC-ec1 (PDB 1OTU [21]), has been considered to represent a more open conformation; however, the permeation pathway in this conformation remains constricted [41]. In CLC-2, the *O* macrostate shows a GLU_*ex*_ conformation completely different from the “up” conformation in the 1OTU structure (Fig. 6). The GLU_*ex*_ conformation seen in the *U* state is not similar to any known structures.

**Fig 5.**
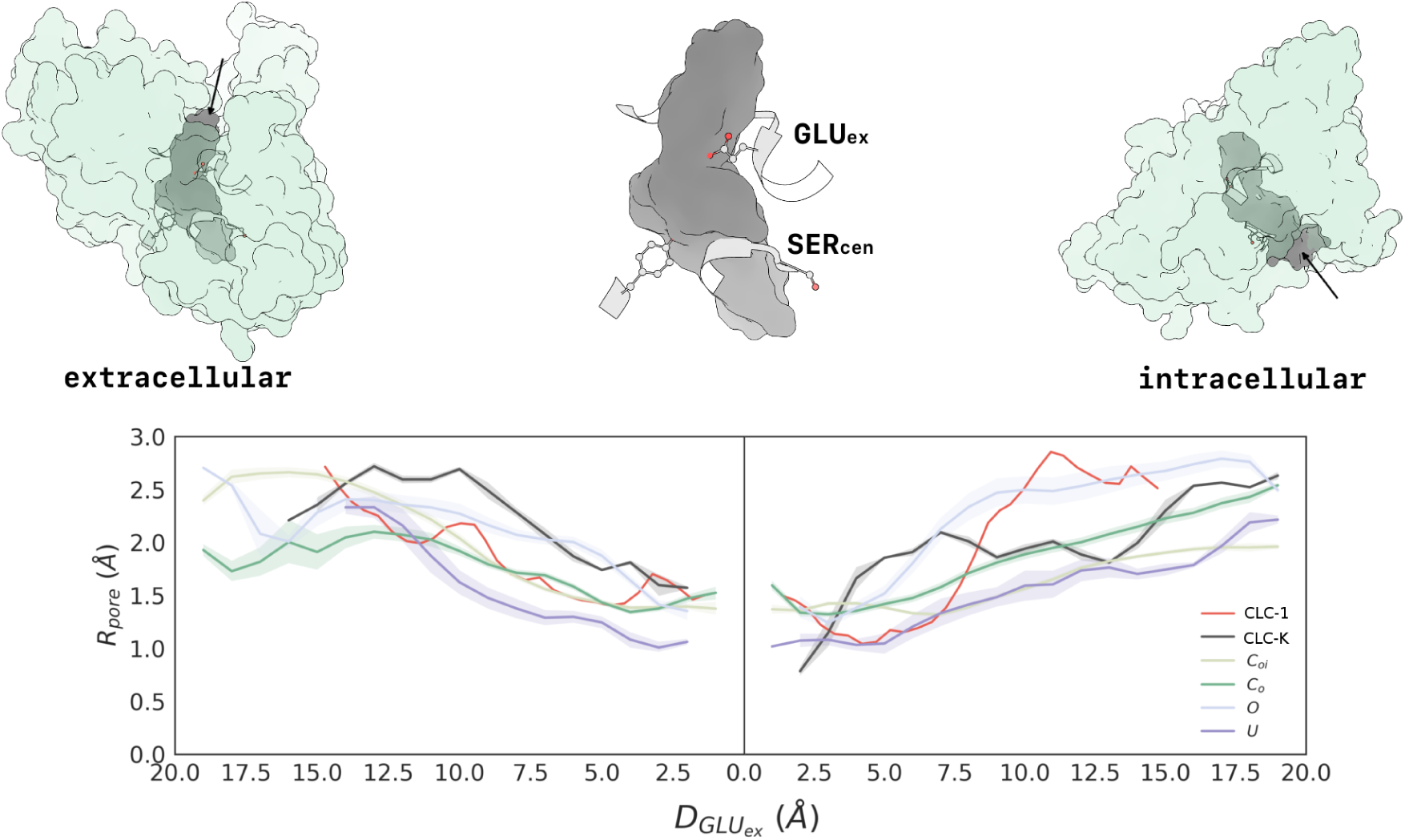
The radius of the ion conduction pathway demonstrates a strong dependence on macrostate. Distance along the pathway (x-axis) is given with respect to the GLU_*ex*_ residue, for each macrostate, and with respect to the VAL mutation for CLC-K (5TQQ, subunit A). The plot on the left illustrates the pore radius from the extracellular side of the protein, descending into the pore to GLU_*ex*_, while the plot on the right illustrates the pore radius from GLU_*ex*_ to the intracellular side of the protein. The structures shown at top depict the volume of the *C*_*o*_ ion conduction pathway from multiple perspectives. All macrostates are relatively open near the extracellular mouth of the protein but narrow in distinct ways to the bottleneck located at GLU_*ex*_. The *O* state is the most open in the region just above GLU_*ex*_, and *U* is the most narrow. Additionally, the minimum radius of the bottleneck is approximately the same for all states except *U*, which is most constricted. Moving toward the intracellular opening from GLU_*ex*_, *O* is once again the most voluminous, while *U* is the most constricted. The pore radius of the CLC-K channel (5TQQ, depicted in black) is wider than the closed-state models of CLC-2 but not as wide overall as the *O* state CLC-2 model. The pore radius of CLC-1 (6COY, depicted in red) has similar dimensions to our *C*_*o*_ and *U* states towards near either side of GLU_*ex*_ but then widens towards the intracellular region along both intracellular pathwas, displaying a radius comparable to our *O* state. Detailed images of the ion conduction pathway and bottleneck for the *O* state, CLC-K, and CLC-1 are shown in the supplementary information, **??, ??**, and **??**, respectively.

**Fig 6.**
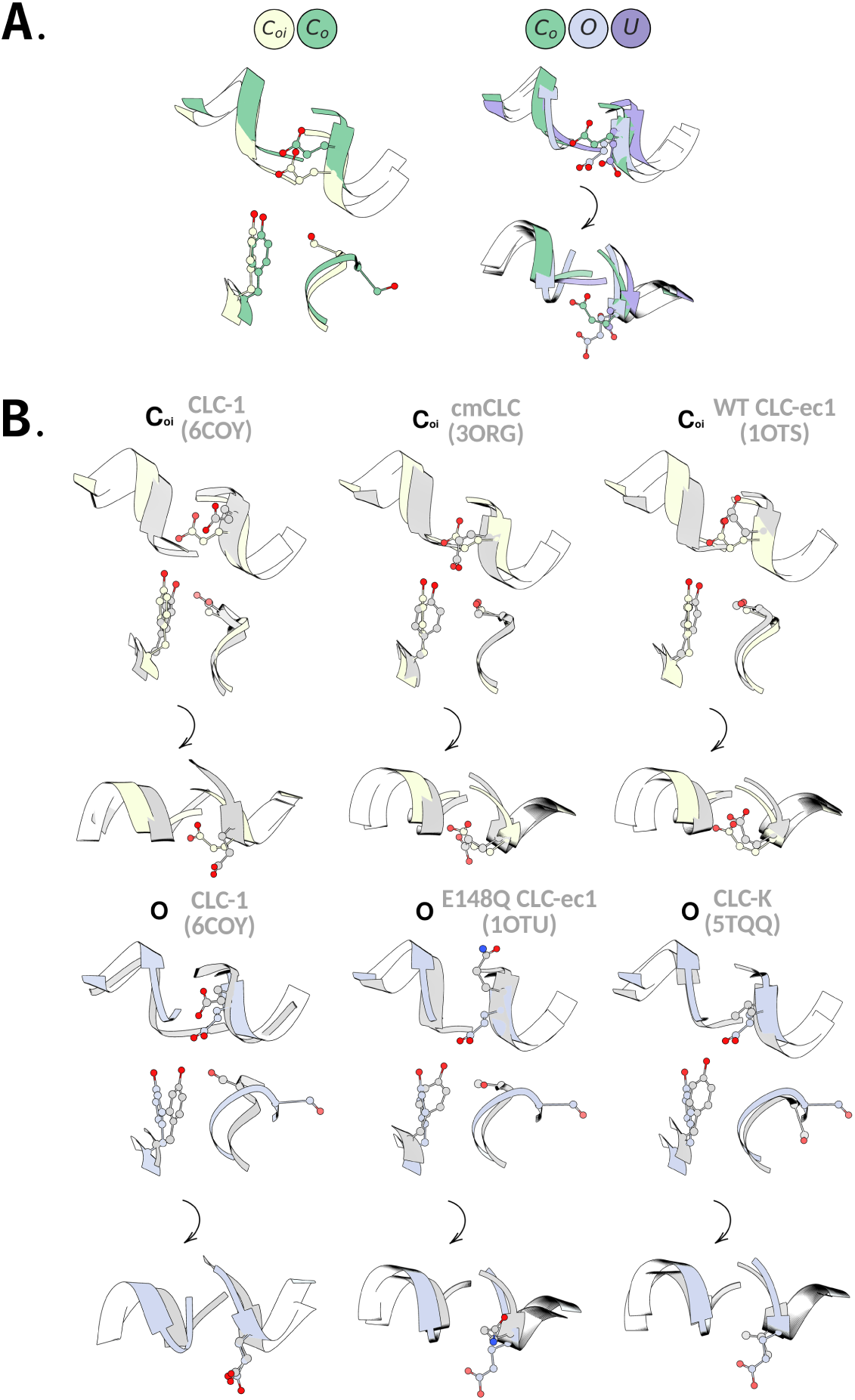
Structural overlays of SF conformations. **A**. Overlay of the four macrostates observed in this study. Left: The S168-G169-I170 backbone conformation distinguishes the *C*_*oi*_ and *C*_*o*_ states. Right: The GLU_*ex*_ side chain conformation distinguishes the *C*_*o*_, *O*, and *U* states. These rotamers are depicted from the plane of the membrane (upper), and also from the extracellular plane (lower). The rotation of SF orientation is signified by the black arrow. **B**. Comparison of states *C*_*oi*_ (top) and *O* (bottom) to relevant structures in the PDB. As in A (right panel), these comparisons are shown from the plane of the membrane (upper), and also from the extracellular plane (lower). The stable *C*_*oi*_ state adopts a GLU_*ex*_ conformation different from that in the template structure 3ORG (cmCLC) and more similar to that observed in most transporter CLC structures in the PDB, as represented by 1OTS (CLC-ec1). The *O* state adopts a GLU_*ex*_ conformation not seen in any known structure, including 1OTU, where GLU_*ex*_ is mutated to GLN, and CLC-1, both of which have been proposed to represent open states. At the intracellular side, the conformation of the S168-G169-I170 backbone of the *O* state resembles that of the 5TQQ (CLC-K) structure. Movies further illustrating overlays of the *C*_*oi*_ and *O* states with the CLC-1 and CLC-K structures are linked to in the SI.

While states *C*_*oi*_, *C*_*o*_, and *O* display bottlenecks of ∼1.5 Å in radius (narrower than the ∼1.8 Å radius of a Cl^−^ ion), the extent of these bottlenecks along the length of the channel pore varies significantly between the conformations (Fig. 5). For *C*_*oi*_ and *C*_*o*_, the pore is constricted to less than 1.8 Å for a ∼19 Å stretch, whereas for state *O*, the pore is constricted for less than half that distance (Fig. 5). Comparing *C*_*oi*_ and *C*_*o*_, the most striking difference is along the intracellular pathway, which widens sufficiently to accommodate chloride in *C*_*o*_. This finding aligns with our initial observation that these states are distinguished by the conformational state of the inner gate, while the outer gate remains closed. The structural difference between these two states is depicted in the overlay of representative conformations shown in Fig. 1 (left panel).

Across both the extracellular and intracellular region, the *O* state is the most voluminous. This finding aligns with our observation that both inner- and outer-gate residues (GLU_*ex*_ and SER_*cen*_) have rotated away from S_*cen*_. The fact that a narrow constriction persists within the putative “open” state (the most open of the set of macrostates) suggests that further conformational change may be involved in reaching a fully open state. Such an additional conformational change (to widen the pore following the rotation of GLU_*ex*_) has been proposed based on experimental studies [41, 79, 94]. For both extracellular and intracellular pathways, the *U* state is the most constricted, despite the fact that the inner-gate SER residue is rotated to the “open” position. The GLU_*ex*_ residue is in a unique rotameric state, rotated away from SER_*cen*_, but this rotation does not increase the volume of the extracellular region of the channel, in part due to the distortion of helix N in this state (see Fig. 2). The constricted nature of the *U* state suggests that it is a non-conductive (closed) channel. That the *U* state is only accessible when GLU_*ex*_ is protonated (neutral) is in agreement with experimental observation that low extracellular pH induces channel closure [22, 93, 95] (though it is known that an extracellular histidine is also involved [96]). Additional analysis of the conformation of the intracellular region can be found in **??**, which shows a Ramachandran plot of the S168-G169 backbone for the *O* macrostate, *C*_*oi*_ macrostate, cmCLC (3ORG) structure, and CLC-K (5TQQ) structure.

Since Cl^−^ binding, unbinding, and translocation play central roles in CLC channel gating [108], we calculated binding-site occupancies for each of the macrostate conformations. The MSM was used to randomly sample 1000 conformations for each state. For each sample, binding site occupancy was computed using the UCSF Chimera occupancy analysis tool [104].

The results, reported in Table 2, reveal that spontaneaous Cl^−^ binding and unbinding occur in a highly macrostate-dependent manner during the simulations. Most strikingly, the high Cl^−^ occupancy of S_*ext*_ binding site in the *O* macrostate is consistent with the notion that filling of S_*ext*_ with Cl^−^ is a driving force for keeping the gate open [14, 17, 22, 82]. Surprisingly, we found little occupancy of the S_*cen*_ site in any state, which in contrary to the observation of Cl^−^ in this site in most crystal structures [108] and to experimental evidence indicating that multi-ion occupancy facilitates CLC-2 opening [73]. Future simulations using alternate ions such as Br^−^ or SCN^−^, which greatly increase CLC-2 opening at zero millivolts [73], and evaluating effects of transmembrane voltage, will be valuable for further illuminating the details of ion-driven gating in CLC-2.

**Table 2.** Occupation statistics of each ion binding site for each conformational state. Values denote the empirical frequency.

|  | *S<sub>ext</sub>* | *S<sub>cen</sub>* | *S<sub>int</sub>* |
| --- | --- | --- | --- |
| *C<sub>oi</sub>* | 0.01 | 0.00 | 0.02 |
| *C<sub>o</sub>* | 0.07 | 0.01 | 0.02 |
| *O* | 0.93 | 0.04 | 0.04 |
| *U* | 0.11 | 0.04 | 0.04 |

The identification of four CLC-2 macrostates with differing pore radii raises important questions about gating and permeation in all CLC channels. All four CLC-2 macrostates have pore dimensions that are significantly wider than those seen in the CLC transporters, which exhibit pinch points at the GLU_*ex*_ and SER_*cen*_ sites. In the transporters, the intracellular (SER_*cen*_) pinch point is thought to act as a “kinetic barrier” that is critical for maintaining strict coupling between Cl^−^ and H^+^ antiport in the CLC transporters [87]. In the CLC-K and CLC-1 channel structures, the widening of this pinch point (reduction of the kinetic barrier) is thought to facilitate rapid Cl^−^ conduction. However, both cryo-EM structures exhibit constrictions that are narrower than the CLC-2 *O* state (Fig. 5). In CLC-1, this relative constriction occurs at the intracellular vestibule where our CLC-2 model predicts a conformational flip of SER_*cen*_, analogous to that seen in the CLC-K structure (Fig. 6B). In CLC-K, this flip is thought to occur due to a sterically bulky tyrosine (Y425) that would clash with SER_*cen*_ in the canonical protein conformation. In CLC-1, this residue is much smaller (glycine), and the movement of SER_*cen*_ thus appears unnecessary [33]. However, our observation that SER_*cen*_ can flip in CLC-2 (which has an alanine at the Y425 position) suggests that this conformation may not be unique to CLC-K and might represent a more general mechanism of CLC gating.

Comparison of the CLC-K and CLC-1 structures to the extracellular pore of our CLC-2 macrostates is also interesting. In state *O*, the extracellular pore is most similar to that of CLC-K. Structural overlays show that GLU_*ex*_ of state *O* closely overlays with the corresponding VAL in the CLC-K structure, which points away from the channel pore (Fig. 6B, bottom right). In contrast, the CLC-1 extracellular pore radius is most similar to that of the CLC-2 *C*_*o*_ state (though it widens to a much greater extent (∼ 2.5 Å) at the initial segment). Structural overlays show that GLU_*ex*_ in CLC-1 is distinct from all of our states (though it is more similar to the *O* state than to the closed states) (Fig. 6B). Perhaps the different GLU_*ex*_ conformations in CLC-1 and CLC-2 are partially responsible for their differences in voltage- and H^+^-dependent gating.

Regardless of the subtle differences between homologs, our results agree with the general conclusion that CLC channels (as opposed to transporters) have lowered kinetic barriers to ion permeation as a result of pore widening at the inner and outer gates. Our modeling, together with recent channel structures and experimental data, provides a framework for addressing the many lingering questions about the molecular mechanisms of CLC gating. Why, for instance, do CLC-1 and CLC-2 exhibit opposite voltage dependencies? What is the structural basis connecting Ca^2+^-binding to channel opening in the CLC-K channels? Is the second intracellular anion pathway observed in the CLC-1 structure [33] relevant in CLC-2, the most closely related homolog? Answers to these and many other questions require further analysis of channel conformational dynamics, including simulations that probe specific Cl^−^/protein interactions, given the key role that the Cl^−^ ion plays in CLC channel gating.

### Analysis of reaction coordinates

The structures identified for the major conformational states were arranged into a schematic network mechanism, given in Fig. 7. This network diagram summarizes the results presented in **??**. We observe that there is a dominant motion from the *C*_*oi*_ to *C*_*o*_ state, which involves the rotation of the S168-G169-I170 backbone. Following this motion, the GLU_*ex*_ side chain may rotate away from the ion conduction pathway. This GLU_*ex*_ flip has a low equilibrium population under these conditions (0 mV), such that return to the resting GLU_*ex*_ conformation is favored. The *O* rotamer possibly activates the channel to the full conducting state, while the protonation-dependent *U* rotamer constricts the conduction pathway to a non-conducting state.

**Fig 7.**
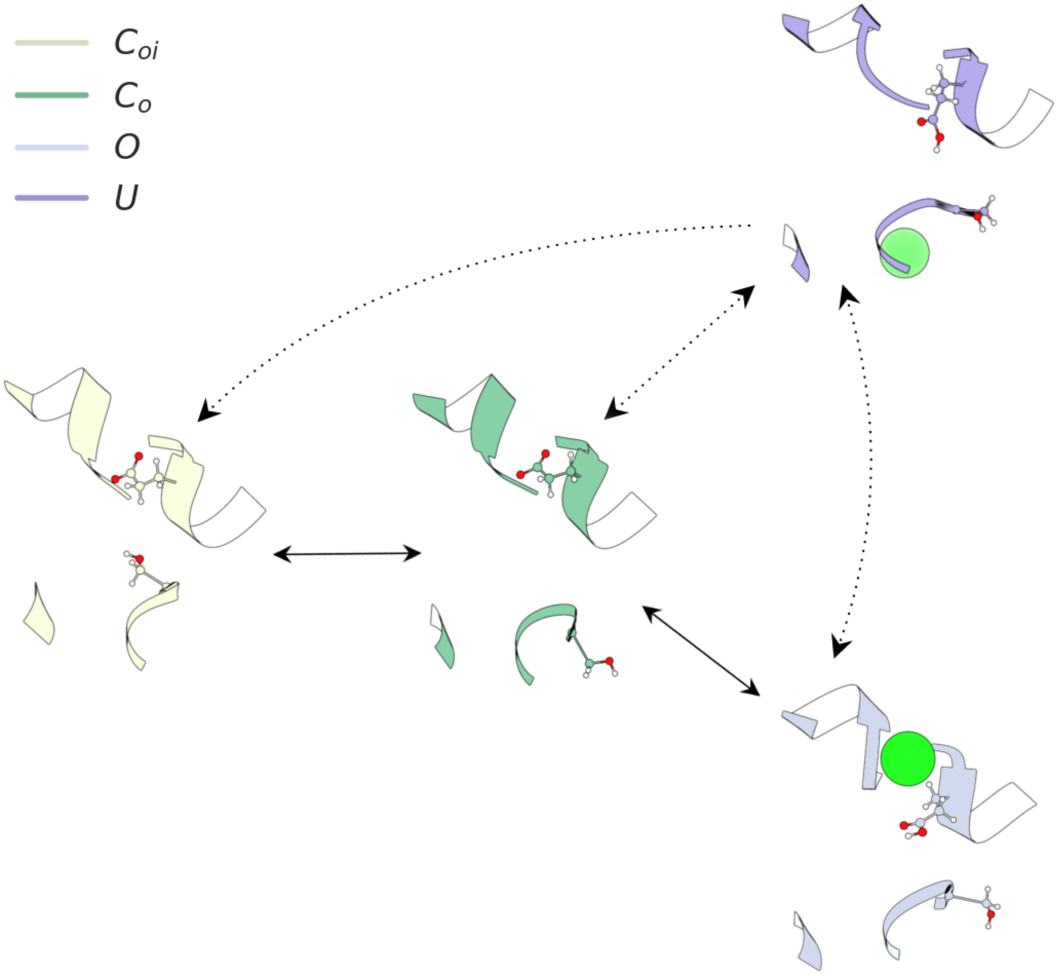
Model of CLC-2 activation at 0 mV and symmetric Cl^−^. Dashed rates signify transitions only accessible when GLU_*ex*_ is protonated. The S168-G169-I170 backbone may exist in two possible conformational states. One points toward the ion conduction pathway, the other has the S168 side chain intracellularly solvent-exposed. The GLU_*ex*_ residue side chain may exist in three possible conformational states. In the closed state, GLU_*ex*_ rests directly in the S_*ext*_ ion binding site, obstructing the ion conduction pathway. When GLU_*ex*_ is protonated, or an ion permeates close to the residue, it flips outward. In the *O* (putative open) state, GLU_*ex*_ moves to the side, opening up space for a chloride ion. When GLU_*ex*_ is protonated, the SF can adopt a third conformation (the *U* state), which constricts the permeation pathway to a non-conducting state.

### Summary and conclusions

We have applied computational modeling methodologies to predict the conformational dynamics of CLC-2 activation. Four major conformational states were identified and characterized with respect to thermodynamics, kinetics, and structure. Of the major states, two were found to be highly stable, and two to be less stable but still detectably populated under the simulation conditions (150 mM Cl^−^, 0 mV). Progression through these states was found to follow a particular sequence under the simulation conditions. First, from the *C*_*oi*_, or closed state, there is flip of the S168-G169-I170 backbone. This flip occurs via an intracellular rotation and results in a volume increase in the inner regions of the ion conduction pathway. Once this occurs, the GLU_*ex*_ side chain is able to flip. Our model suggests that intracellular gate opening (mediated by the intracellular rotation of SER_*cen*_) must occur prior to opening of the extracellular gate (rotation of GLU_*ex*_). Such a mechanism is consistent with the picture emerging from mechanistic electrophysiological studies, which indicate that intracellular Cl^−^ is the main factor driving CLC-2 channel opening [73]. The intracellular rotation of the S168-G169-I170 backbone that we observe would allow intracellular chloride to access the inner channel pore and facilitate opening of the outer glutamate gate by forcing GLU_*ex*_ into its open conformation as chloride traverses the membrane.

Our studies of GLU_*ex*_ protonation reveal consistencies with the existing literature, and they suggest potential areas of new investigation. Our observation that protonation of GLU_*ex*_ is not required for opening is consistent with functional studies of CLC-2 [73], as is our observation that protonation of GLU_*ex*_ decreases the rate of transition from open to closed conformations [74]. Our observation that protonation of GLU_*ex*_ introduces the ability to adopt an alternative rotamer orientation suggests a potential link between CLC-2 inhibition by extracellular protons (which also involves an extracellular histidine residue [96]). Given the complex relationship between H^+^, Cl^−^, and voltage in regulating CLC-2 gating, many questions remain. This CLC-2 activation model, together with future simulations to explore Cl^−^/protein interactions in detail, will complement current and future efforts toward furthering the biophysical understanding of CLC proteins.

## Methods

### Molecular dynamics

The initial structures consisted of a CLC-2 homology model built using the software Modeller [97]. The rat CLC-2 monomer sequence was mapped onto a monomer of the 3ORG structure. Unresolved regions of the template structure were refined via Modeller. The membrane system was assembled via CHARMMGUI [98]. The simulated system consisted of the homology model structure, 2 leaflets of 64 3:1 POPE:POPG lipid molecules, explicit water, and 0.15 M K^+^Cl^−^. The TIP3P [99], lipid14 [100] and AMBER14SB [101] forcefields were used. Protonation of the GLU_*ex*_ residue was accomplished using CHARMMGUI. The protonation of all other residues was given by the expected protonation state for equilibrium physiological pH. Equilibration was done using AMBER 14*, with restraints on the protein and lipids, which were slowly released as the system was heated gradually from 0 to 300 K. An additional 100 ns simulation was run, from which initial system configurations were sampled. The AMBER prmtop and inpcrd files of these structures were used to set up OpenMM [102] simulations in the NPT ensemble. From these initial structures, 100 unique velocity distributions were assigned. Simulations were performed at 0 mV. This is due to the novelty of applying membrane potentials in simulation and concern about the accurate response of MD force fields to applied external electric fields. The distributed computing platform folding@home [103] was then used to perform 600 *µ*s of aggregate MD simulation. Approximately 1000 parallel simulations were run, for a length averaging 600 ns. The UCSF Chimera program [104] was used to visualize the MD trajectories and generate figures.

### Markov state modeling

MSM analysis was accomplished using MSMBuilder 3.7 [105]. The MD datasets were featurized using the signed *ϕ, ψ, χ*_1_, and *χ*_2_ dihedral angles for all residues in the CLC-2 SF. The SF residue numbers are as follows (using the rCLC-2 primary sequence numbering): 167, 168, 169, 170, 171, 209, 210, 211, 212, 213, 463, 464, 465, 466, 467, 468, 559. Note that the MSM was constructed strictly over protein conformation. The presence of lipid, water, and chloride in the simulations means that the impact of these elements on CLC-2 protein conformations is implicitly accounted for. Detailed analyses of specific interactions between lipid, water, chloride, and protein, which will require methodological development, stand as an exciting avenue for future work. The featurized datasets were transformed using kinetic-mapping of all tICA tICs. The transformed data was clustered into 324 microstates via the mini-batch kmeans method. The same tICA model and state decomposition was used for both datasets. 100 bootstrap MSMs were constructed (with a lag-time of 28.8 ns, see **??**), from which the maximum likelihood estimate model has been highlighted. The state equilibrium populations were derived from the first eigenvector of the MSM transition matrix. Error bars for the free energy estimates were calculated using all MSM models. Macrostates were determined using the PCCA+ algorithm. A continuous time rate-matrix MSM was used to estimate the kinetic rates of macrostate conversion. The Caver 3.0 software package [106] was used to calculate the pore radii for each macrostate.

## Acknowledgements

We acknowledge Professor Justin Du Bois for helpful discussion. We acknowledge the Folding@home donors who contributed to this project (PROJ9625-9643).

## References

1. Jentsch TJ. Discovery of CLC transport proteins: cloning, structure, function and pathophysiology. The Journal of physiology. 2015;593(18):4091–4109.

2. Stauber T, Weinert S, Jentsch TJ. Cell biology and physiology of CLC chloride channels and transporters. Comprehensive Physiology. 2012;.

3. Stölting G, Fischer M, Fahlke C. CLC channel function and dysfunction in health and disease. Frontiers in physiology. 2014;5.

4. Poroca DR, Pelis RM, Chappe VM. CLC Channels and Transporters: Structure, Physiological Functions, and Implications in Human Chloride Channelopathies. Frontiers in pharmacology. 2017;8.

5. De Angeli A, Monachello D, Ephritikhine G, Frachisse JM, Thomine S, Gambale F, et al. CLC-mediated anion transport in plant cells. Philosophical Transactions of the Royal Society of London B: Biological Sciences. 2009;364(1514):195–201.

6. Iyer R, Iverson TM, Accardi A, Miller C. A biological role for prokaryotic CLC chloride channels. Nature. 2002;419(6908):715–718.

7. Fong P. Twenty-five years of CLC chloride transport proteins. The Journal of physiology. 2015;593(18):4083–4084.

8. Miller C. CLC chloride channels viewed through a transporter lens. Nature. 2006;440(7083):484–489.

9. Accardi A. Structure and gating of CLC channels and exchangers. The Journal of physiology. 2015;593(18):4129–4138.

10. Middleton RE, Pheasant DJ, Miller C. Homodimeric architecture of a CLC-type chloride ion channel. Nature. 1996;383(6598):337.

11. Ludewig U, Pusch M, Jentsch TJ. Two physically distinct pores in the dimeric CLC-0 chloride channel. Nature. 1996;383(6598):340–343.

12. Maduke M, Miller C, Mindell JA. A decade of CLC chloride channels: structure, mechanism, and many unsettled questions. Annual review of biophysics and biomolecular structure. 2000;29(1):411–438.

13. Dutzler R, Campbell EB, Cadene M, Chait BT, MacKinnon R. X-ray structure of a CLC chloride channel at 3.0 Å reveals the molecular basis of anion selectivity. Nature. 2002;415(6869):287–294.

14. Dutzler R. Structural basis for ion conduction and gating in ClC chloride channels. FEBS letters. 2004;564(3):229–233.

15. Lin CW, Chen TY. Probing the pore of CLC-0 by substituted cysteine accessibility method using methane thiosulfonate reagents. The Journal of general physiology. 2003;122(2):147–159.

16. Engh AM, Maduke M. Cysteine accessibility in CLC-0 supports conservation of the CLC intracellular vestibule. The Journal of general physiology. 2005;125(6):601–617.

17. Engh AM, Faraldo-Gomez Jose D, Maduke M. The mechanism of fast-gate opening in ClC-0. The Journal of general physiology. 2007;130(4):335–349.

18. Matulef K, Maduke M. The CLC ‘chloride channel’ family: revelations from prokaryotes. Molecular membrane biology. 2007;24(5-6):342–350.

19. Estévez R, Schroeder BC, Accardi A, Jentsch TJ, Pusch M. Conservation of chloride channel structure revealed by an inhibitor binding site in CLC-1. Neuron. 2003;38(1):47–59.

20. Lísal J, Maduke M. The CLC-0 chloride channel is a ‘broken’ Cl^−^/H^+^ antiporter. Nature structural & molecular biology. 2008;15(8):805–810.

21. Dutzler R, Campbell EB, MacKinnon R. Gating the selectivity filter in CLC chloride channels. Science. 2003;300(5616):108–112.

22. Niemeyer MI, Cid LP, Zúñiga L, Catalán M, Sepúlveda FV. A conserved pore-lining glutamate as a voltage-and chloride-dependent gate in the ClC-2 chloride channel. The Journal of physiology. 2003;553(3):873–879.

23. Vien M, Basilio D, Leisle L, Accardi A. Probing the conformation of a conserved glutamic acid within the Cl-pathway of a CLC H+/Cl-exchanger. The Journal of General Physiology. 2017; p. jgp–201611682.

24. Phillips S, Brammer AE, Rodriguez L, Lim HH, Stary-Weinzinger A, Matulef K. Surprises from an unusual CLC homolog. Biophysical journal. 2012;103(9):L44–L46.

25. Picollo A, Malvezzi M, Accardi A. Proton block of the CLC-5 Cl-/H+ exchanger. The Journal of general physiology. 2010;135(6):653–659.

26. Costa A, Gutla PVK, Boccaccio A, Scholz-Starke J, Festa M, Basso B, et al. The Arabidopsis central vacuole as an expression system for intracellular transporters: functional characterization of the Cl-/H+ exchanger CLC-7. The Journal of physiology. 2012;590(15):3421–3430.

27. Bell SP, Curran PK, Choi S, Mindell JA. Site-directed fluorescence studies of a prokaryotic ClC antiporter. Biochemistry. 2006;45(22):6773–6782.

28. de Santiago JA, Nehrke K, Arreola J. Quantitative analysis of the voltage-dependent gating of mouse parotid ClC-2 chloride channel. The Journal of general physiology. 2005;126(6):591–603.

29. Basilio D, Noack K, Picollo A, Accardi A. Conformational changes required for H^+^/Cl^−^exchange mediated by a CLC transporter. Nature structural & molecular biology. 2014;21(5):456–463.

30. Jayaram H, Accardi A, Wu F, Williams C, Miller C. Ion permeation through a Cl^−^selective channel designed from a CLC Cl^−^/H^+^ exchanger. Proceedings of the National Academy of Sciences. 2008;105(32):11194–11199.

31. Elvington SM, Liu CW, Maduke MC. Substrate-driven conformational changes in CLC-ec1 observed by fluorine NMR. The EMBO journal. 2009;28(20):3090–3102.

32. Park E, Campbell EB, MacKinnon R. Structure of a CLC chloride ion channel by cryo-electron microscopy. Nature. 2017;541(7638):500–505.

33. Park E, MacKinnon R. Structure of the CLC-1 chloride channel from Homo sapiens. Elife. 2018;7:e36629.

34. Pusch M, Ludewig U, Rehfeldt A, Jentsch TJ. Gating of the voltage-dependent chloride channel CLC-0 by the permeant anion. Nature. 1995;373(6514):527–531.

35. Chen TY, Miller C. Nonequilibrium gating and voltage dependence of the CLC-0 Cl^−^channel. The Journal of General Physiology. 1996;108(4):237–250.

36. Lísal J, Maduke M. Proton-coupled gating in chloride channels. Philosophical Transactions of the Royal Society of London B: Biological Sciences. 2009;364(1514):181–187.

37. Accardi A, Picollo A. CLC channels and transporters: proteins with borderline personalities. Biochimica et Biophysica Acta (BBA)-Biomembranes. 2010;1798(8):1457–1464.

38. Pusch M. Structural insights into chloride and proton-mediated gating of CLC chloride channels. Biochemistry. 2004;43(5):1135–1144.

39. Lee S, Mayes HB, Swanson JM, Voth GA. The Origin of Coupled Chloride and Proton Transport in a Cl^−^/H^+^ Antiporter. J Am Chem Soc. 2016;138(45):14923–14930.

40. Jiang T, Han W, Maduke M, Tajkhorshid E. Molecular Basis for Differential Anion Binding and Proton Coupling in the Cl^−^/H^+^ Exchanger ClC-ec1. Journal of the American Chemical Society. 2016;138(9):3066–3075.

41. Khantwal CM, Abraham SJ, Han W, Jiang T, Chavan TS, Cheng RC, et al. Revealing an outward-facing open conformational state in a CLC Cl^−^/H^+^ exchange transporter. eLife. 2016;5:e11189.

42. Cheng MH, Coalson RD. Molecular dynamics investigation of Cl^−^and water transport through a eukaryotic CLC transporter. Biophysical journal. 2012;102(6):1363–1371.

43. Ko YJ, Jo WH. Chloride ion conduction without water coordination in the pore of ClC protein. Journal of computational chemistry. 2010;31(3):603–611.

44. Bisset D, Corry B, Chung SH. The fast gating mechanism in ClC-0 channels. Biophysical journal. 2005;89(1):179–186.

45. Bostick DL, Berkowitz ML. Exterior site occupancy infers chloride-induced proton gating in a prokaryotic homolog of the ClC chloride channel. Biophysical journal. 2004;87(3):1686–1696.

46. Cohen J, Schulten K. Mechanism of anionic conduction across ClC. Biophysical journal. 2004;86(2):836–845.

47. Picollo A, Xu Y, Johner N, Bernèche S, Accardi A. Synergistic substrate binding determines the stoichiometry of transport of a prokaryotic H+/Cl-exchanger. Nature structural & molecular biology. 2012;19(5):525–531.

48. Lee S, Swanson JM, Voth GA. Multiscale simulations reveal key aspects of the proton transport mechanism in the ClC-ec1 antiporter. Biophysical journal. 2016;110(6):1334–1345.

49. Thiemann A, Grunder S, et al. A chloride channel widely expressed in epithelial and non-epithelial cells. Nature. 1992;356(6364):57.

50. Blanz J, Schweizer M, Auberson M, Maier H, Muenscher A, Hübner CA, et al. Leukoencephalopathy upon disruption of the chloride channel CLC-2. Journal of Neuroscience. 2007;27(24):6581–6589.

51. Jeworutzki E, López-Hernández T, Capdevila-Nortes X, Sirisi S, Bengtsson L, Montolio M, et al. GlialCAM, a protein defective in a leukodystrophy, serves as a ClC-2 Cl-channel auxiliary subunit. Neuron. 2012;73(5):951–961.

52. Hoegg-Beiler MB, Sirisi S, Orozco IJ, Ferrer I, Hohensee S, Auberson M, et al. Disrupting MLC1 and GlialCAM and ClC-2 interactions in leukodystrophy entails glial chloride channel dysfunction. Nature communications. 2014;5:3475.

53. Depienne C, Bugiani M, Dupuits C, Galanaud D, Touitou V, Postma N, et al. Brain white matter oedema due to CLC-2 chloride channel deficiency: an observational analytical study. The Lancet Neurology. 2013;12(7):659–668.

54. Jentsch TJ. From mice to man: chloride transport in leukoencephalopathy. The Lancet Neurology. 2013;12(7):626–628.

55. Ge YX, Liu Y, Tang HY, Liu XG, Wang X. ClC-2 contributes to tonic inhibition mediated by *α*5 subunit-containing GABA A receptor in experimental temporal lobe epilepsy. Neuroscience. 2011;186:120–127.

56. Madison DV, Malenka RC, Nicoll RA. Phorbol esters block a voltage-sensitive chloride current in hippocampal pyramidal cells. Nature. 1986;321(6071):695–697.

57. Földy C, Lee SH, Morgan RJ, Soltesz I. Regulation of fast-spiking basket cell synapses by the chloride channel ClC-2. Nature neuroscience. 2010;13(9):1047–1049.

58. Clark S, Jordt SE, Jentsch TJ, Mathie A. Characterization of the hyperpolarization-activated chloride current in dissociated rat sympathetic neurons. The Journal of Physiology. 1998;506(3):665–678.

59. Staley K, Smith R, Schaack J, Wilcox C, Jentsch TJ. Alteration of GABA A receptor function following gene transfer of the CLC-2 chloride channel. Neuron. 1996;17(3):543–551.

60. Rinke I, Artmann J, Stein V. ClC-2 voltage-gated channels constitute part of the background conductance and assist chloride extrusion. Journal of Neuroscience. 2010;30(13):4776–4786.

61. Kleefuss-Lie A, Friedl W, Cichon S, Haug K, Warnstedt M, Alekov A, et al. CLCN2 variants in idiopathic generalized epilepsy. Nature genetics. 2009;41(9):954–955.

62. D’agostino D, Bertelli M, Gallo S, Cecchin S, Albiero E, Garofalo PG, et al. Mutations and polymorphisms of the CLCN2 gene in idiopathic epilepsy. Neurology. 2004;63(8):1500–1502.

63. Cortez M, Li C, Whitehead S, Dhani S, D’Antonio C, Huan L, et al. Disruption of ClC-2 expression is associated with progressive neurodegeneration in aging mice. Neuroscience. 2010;167(1):154–162.

64. Niemeyer MI, Cid LP, Sepúlveda FV, Blanz J, Auberson M, Jentsch TJ. No evidence for a role of CLCN2 variants in idiopathic generalized epilepsy. Nature genetics. 2010;42(1):3–3.

65. Ratté S, Prescott SA. ClC-2 channels regulate neuronal excitability, not intracellular chloride levels. Journal of Neuroscience. 2011;31(44):15838–15843.

66. Pande VS, Beauchamp K, Bowman GR. Everything you wanted to know about Markov State Models but were afraid to ask. Methods. 2010;52(1):99 – 105. doi:10.1016/j.ymeth.2010.06.002.

67. Schwantes CR, McGibbon RT, Pande VS. Perspective: Markov models for long-timescale biomolecular dynamics. J Chem Phys. 2014;141(9):090901. doi:10.1063/1.4895044.

68. Chodera JD, Noé F. Markov state models of biomolecular conformational dynamics. Curr Opin Struct Biol. 2014;25:135–144. doi:10.1016/j.sbi.2014.04.002.

69. Bowman GR, Pande VS, Noé F. An Introduction to Markov State Models and Their Application to Long Timescale Molecular Simulation. vol. 797. Springer; 2014.

70. Schwantes CR, Pande VS. Improvements in Markov state model construction reveal many non-native interactions in the folding of NTL9. Journal of chemical theory and computation. 2013;9(4):2000–2009.

71. Pérez-Hernández G, Paul F, Giorgino T, De Fabritiis G, Noé F. Identification of slow molecular order parameters for Markov model construction. The Journal of chemical physics. 2013;139(1):07B604 1.

72. Naritomi Y, Fuchigami S. Slow dynamics in protein fluctuations revealed by time-structure based independent component analysis: The case of domain motions. The Journal of chemical physics. 2011;134(6):02B617.

73. De Jesús-Pérez JJ, Castro-Chong A, Shieh RC, Hernández-Carballo CY, De Santiago-Castillo JA, Arreola J. Gating the glutamate gate of CLC-2 chloride channel by pore occupancy. The Journal of general physiology. 2016;147(1):25–37.

74. Sánchez-Rodríguez JE, Santiago-Castillo D, José A, Contreras-Vite JA, Nieto-Delgado PG, Castro-Chong A, et al. Sequential interaction of chloride and proton ions with the fast gate steer the voltage-dependent gating in ClC-2 chloride channels. The Journal of physiology. 2012;590(17):4239–4253.

75. Yu Y, Tsai MF, Yu WP, Chen TY. Modulation of the slow/common gating of CLC channels by intracellular cadmium. The Journal of general physiology. 2015;146(6):495–508.

76. Duffield M, Rychkov G, Bretag A, Roberts M. Involvement of helices at the dimer interface in ClC-1 common gating. The Journal of general physiology. 2003;121(2):149–161.

77. Pusch M, Ludewig U, Jentsch TJ. Temperature dependence of fast and slow gating relaxations of ClC-0 chloride channels. The Journal of general physiology. 1997;109(1):105–116.

78. Ludewig U, Pusch M, Jentsch TJ. Independent gating of single pores in CLC-0 chloride channels. Biophysical Journal. 1997;73(2):789–797.

79. Accardi A, Pusch M. Conformational changes in the pore of CLC-0. The Journal of general physiology. 2003;122(3):277–294.

80. Traverso S, Zifarelli G, Aiello R, Pusch M. Proton sensing of CLC-0 mutant E166D. The Journal of general physiology. 2006;127(1):51–66.

81. Zifarelli G, Pusch M. The role of protons in fast and slow gating of the Torpedo chloride channel ClC-0. European Biophysics Journal. 2010;39(6):869–875.

82. Chen TY. Coupling gating with ion permeation in ClC channels. Science Signaling. 2003;2003(188):pe23–pe23.

83. Chen TY, Chen MF, Lin CW. Electrostatic control and chloride regulation of the fast gating of ClC-0 chloride channels. The Journal of general physiology. 2003;122(5):641–651.

84. Sánchez-Rodríguez JE, Santiago-Castillo D, José A, Arreola J. Permeant anions contribute to voltage dependence of ClC-2 chloride channel by interacting with the protopore gate. The Journal of physiology. 2010;588(14):2545–2556.

85. Pusch M, Jordt SE, Stein V, Jentsch TJ. Chloride dependence of hyperpolarization-activated chloride channel gates. The Journal of Physiology. 1999;515(2):341–353.

86. Zifarelli G, Murgia AR, Soliani P, Pusch M. Intracellular proton regulation of ClC-0. The Journal of general physiology. 2008;132(1):185–198.

87. Feng L, Campbell EB, Hsiung Y, MacKinnon R. Structure of a eukaryotic CLC transporter defines an intermediate state in the transport cycle. Science. 2010;330(6004):635–641.

88. Crooks GE, Hon G, Chandonia JM, Brenner SE. WebLogo: a sequence logo generator. Genome research. 2004;14(6):1188–1190.

89. Schneider TD, Stephens RM. Sequence logos: a new way to display consensus sequences. Nucleic acids research. 1990;18(20):6097–6100.

90. Olivella M, Gonzalez A, Pardo L, Deupi X. Relation between sequence and structure in membrane proteins. Bioinformatics. 2013;29(13):1589–1592.

91. Schwantes CR, Pande VS. Modeling molecular kinetics with tICA and the kernel trick. Journal of chemical theory and computation. 2015;11(2):600–608.

92. Chen MF, Chen TY. Side-chain charge effects and conductance determinants in the pore of ClC-0 chloride channels. The Journal of general physiology. 2003;122(2):133–145.

93. Arreola J, Begenisich T, Melvin JE. Conformation-dependent regulation of inward rectifier chloride channel gating by extracellular protons. The Journal of physiology. 2002;541(1):103–112.

94. Traverso S, Elia L, Pusch M. Gating competence of constitutively open CLC-0 mutants revealed by the interaction with a small organic inhibitor. The Journal of general physiology. 2003;122(3):295–306.

95. Jordt SE, Jentsch TJ. Molecular dissection of gating in the CLC-2 chloride channel. The EMBO Journal. 1997;16(7):1582–1592.

96. Niemeyer MI, Cid LP, Yusef YR, Briones R, Sepúlveda FV. Voltage-dependent and-independent titration of specific residues accounts for complex gating of a ClC chloride channel by extracellular protons. The Journal of physiology. 2009;587(7):1387–1400.

97. Šali A, Blundell TL. Comparative protein modelling by satisfaction of spatial restraints. Journal of molecular biology. 1993;234(3):779–815.

98. Lee J, Cheng X, Swails JM, Yeom MS, Eastman PK, Lemkul JA, et al. CHARMM-GUI Input Generator for NAMD, GROMACS, AMBER, OpenMM, and CHARMM/OpenMM Simulations Using the CHARMM36 Additive Force Field. J Chem Theory Comput. 2016;12(1):405–413. doi:10.1021/acs.jctc.5b00935.

99. Jorgensen WL, Chandrasekhar J, Madura JD, Impey RW, Klein ML. Comparison of simple potential functions for simulating liquid water. The Journal of chemical physics. 1983;79(2):926–935.

100. Dickson CJ, Madej BD, Skjevik ÅA, Betz RM, Teigen K, Gould IR, et al. Lipid14: The Amber Lipid Force Field. J Chem Theory Comput. 2014;10(2):865–879. doi:10.1021/ct4010307.

101. Maier JA, Martinez C, Kasavajhala K, Wickstrom L, Hauser KE, Simmerling C. ff14SB: improving the accuracy of protein side chain and backbone parameters from ff99SB. Journal of chemical theory and computation. 2015;11(8):3696–3713.

102. Eastman P, Pande V. OpenMM: A hardware-independent framework for molecular simulations. Computing in Science & Engineering. 2010;12(4):34–39.

103. Shirts M, Pande VS, et al. Screen savers of the world unite. Computing. 2006;10:43.

104. Pettersen EF, Goddard TD, Huang CC, Couch GS, Greenblatt DM, Meng EC, et al. UCSF Chimera: a visualization system for exploratory research and analysis. Journal of computational chemistry. 2004;25(13):1605–1612.

105. Harrigan MP, Sultan MM, Hernández CX, Husic BE, Eastman P, Schwantes CR, et al. MSMBuilder: Statistical Models for Biomolecular Dynamics. Biophysical journal. 2017;112(1):10–15.

106. Chovancova E, Pavelka A, Benes P, Strnad O, Brezovsky J, Kozlikova B, et al. CAVER 3.0: a tool for the analysis of transport pathways in dynamic protein structures. PLoS computational biology. 2012;8(10):e1002708.

107. Garcia-Olivares, Jennie and Alekov, Alexi and Boroumand, Mohammad Reza and Begemann, Birgit and Hidalgo, Patricia and Fahlke Christoph Gating of human CLC-2 chloride channels and regulation by carboxy-terminal domains. The Journal of physiology. 2008;586(22):5325–5336.

108. Jentsch, Thomas J and Pusch Michael CLC chloride channels and transporters: structure, function, physiology, and disease. American Physiological Society. 2018;98(3):1493–1590.

109. Chadda, Rahul and Krishnamani, Venkatramanan and Mersch, Kacey and Wong, Jason and Brimberry, Marley and Chadda, Ankita and Kolmakova-Partensky, Ludmila and Friedman Larry J and Gelles Jeff and Robertson Janice L The dimerization equilibrium of a ClC Cl-/H+ antiporter in lipid bilayers eLife Sciences Publications Limited. 2016;5:e17438.

110. Varela, Diego and Niemeyer, María Isabel and Cid, L Pablo and Sepúlveda, Francisco V Effect of an N-terminus deletion on voltage-dependent gating of the ClC-2 chloride channel Wiley Online Library. 2002;544(2):363–372.

111. Robertson, Janice L and Kolmakova-Partensky, Ludmila and Miller, Christopher Design, function and structure of a monomeric ClC transporter Nature Publishing Group. 2010;468(7325):844.

112. Laskowski, Roman A and MacArthur, Malcolm W and Moss, David S and Thornton, Janet M PROCHECK: a program to check the stereochemical quality of protein structures International Union of Crystallography. 1993;26(2):468(7325):283–291.

113. Bösl, Michael R and Stein, Valentin and Hübner, Christian and Zdebik, Anselm A and Jordt, Sven-Eric and Mukhopadhyay, Amal K and Davidoff, Michail S and Holstein, Adolf-Friedrich and Jentsch, Thomas J Male germ cells and photoreceptors, both dependent on close cell–cell interactions, degenerate upon ClC-2 Cl-channel disruption The EMBO journal. 2001;20(6):1289–1299.

114. Catalán, Marcelo A and Flores, Carlos A and González–Begne, Mireya and Zhang, Yan and Sepúlveda, Francisco V and Melvin, James E Severe defects in absorptive ion transport in distal colons of mice that lack ClC-2 channels Gastroenterology. 2012;142(2):346–354.

115. Zdebik, Anselm A and Cuffe, John E and Bertog, Marko and Korbmacher, Christoph and Jentsch, Thomas J Additional disruption of the ClC-2 Cl-channel does not exacerbate the cystic fibrosis phenotype of cystic fibrosis transmembrane conductance regulator mouse models Journal of Biological Chemistry. 2004;279(21):22276–22283.

116. Fernandes-Rosa, Fabio L and Daniil, Georgios and Orozco, Ian J and Göppner, Corinna and El Zein, Rami and Jain, Vandana and Boulkroun, Sheerazed and Jeunemaitre, Xavier and Amar, Laurence and Lefebvre, Hervé and others A gain-of-function mutation in the CLCN2 chloride channel gene causes primary aldosteronism Nature genetics. 2018;50(3):355.

